# Annotation of DOM metabolomes with an ultrahigh resolution mass spectrometry molecular formula library

**DOI:** 10.1101/2024.04.30.591926

**Authors:** Nicole R Coffey, Christian Dewey, Kieran Manning, Yuri Corilo, William Kew, Lydia Babcock-Adams, Amy M McKenna, Rhona K Stuart, Rene M Boiteau

**Affiliations:** Department of Earth and Environmental Sciences, University of Minnesota, Minneapolis, Minnesota 55455, USA; Department of Chemistry, University of Minnesota, Minneapolis, Minnesota 55455, USA; College of Earth, Ocean, and Atmospheric Sciences, Oregon State University, Corvallis, Oregon 97331, USA; Environmental Molecular Sciences Laboratory, Pacific Northwest National Laboratory, Richland, Washington 99354, USA; National High Magnetic Field Laboratory, Florida State University, Tallahassee, Florida 32310, USA; Department of Soil and Crop Sciences, Colorado State University, Fort Collins, CO 80523; Physical and Life Sciences Directorate, Lawrence Livermore National Laboratory, Livermore, CA 94550, USA

**Keywords:** algal exometabolome, Dissolved Organic Matter, FT-ICR MS, Metabolomics, Liquid Chromatography Mass Spectrometry, iron stress

## Abstract

Increased accessibility of liquid chromatography mass spectrometry (LC-MS) metabolomics instrumentation and software have expanded their use in studies of dissolved organic matter (DOM) and exometabolites released by microbes. Current strategies to annotate metabolomes generally rely on matching tandem MS/MS spectra to databases of authentic standards. However, spectral matching approaches typically have low annotation rates for DOM. An alternative approach is to annotate molecular formula based on accurate mass and isotopic fine structure measurements that can be obtained from state-of-the-art ultrahigh resolution Fourier Transform Ion Cyclotron Resonance mass spectrometry (FT-ICR-MS), but instrument accessibility for large metabolomic studies is generally limited. Here, we describe a strategy to annotate exometabolomes obtained from lower resolution LC-MS systems by matching metabolomic features to a molecular formula library generated for a representative sample analyzed by LC-21T FT-ICR MS. The molecular formula library approach successfully annotated 53% of exometabolome features of the marine diatom *Phaeodactylum tricornutum* – a nearly ten-fold increase over the 6% annotation rate achieved using a conventional MS/MS approach. There was 94% agreement between assigned formula that were annotated with both approaches, and mass error analysis of the discrepancies suggested that the FT-ICR MS formula assignments were more reliable. Differences in the exometabolome of *P. tricornutum* grown under iron replete and iron limited conditions revealed 668 significant metabolites, including a suite of peptide-like molecules released by *P. tricornutum* in response to iron deficiency. These findings demonstrate the utility of FT-ICR MS formula libraries for extending the accuracy and comprehensiveness of metabolome annotations.

## 1. Introduction

Dissolved organic matter (DOM) represents a major conduit of energy, fixed carbon, and nutrients through the environment. Various factors, including species, growth phase, and nutrient stress, have been shown to alter the chemical characteristics and bioavailability of dissolved organic matter (DOM) released by microorganisms (Grossart and Simon, 2007; Becker et al., 2014; Wear et al., 2015; Tian et al., 2023). However, the specific metabolic processes that govern DOM composition remain a mystery. This limitation stems from the difficulty in separating and identifying individual molecules within the complex mixture of DOM (e.g., Hedges, 2002; Moran et al., 2016; Boiteau et al., 2024). Since the production and consumption of DOM directly influence microbial interactions, such as competition and resource partitioning (e.g., Wang et al., 2022; Bao et al., 2023), our ability to understand microbial community dynamics is also limited.

Liquid chromatography mass spectrometry (LC-MS) based metabolomic analyses hold promise for filling this knowledge gap. These analyses aim to identify and quantify a wide range of small molecules simultaneously across a biosystem. In a standard LC-MS workflow, data analysis software is used to determine the mass, retention time, and abundance of chromatographically resolved ‘features’ that correspond to distinct molecules. To identify molecules, their mass and retention time can be matched to those of authentic standards. When tandem MS/MS capable instruments are used, features are more commonly identified by comparing their fragmentation spectra to databases of MS/MS spectra from authentic standards (e.g., Milman, 2005; Schymanski et al., 2014; Luo et al., 2016, 2023; Folberth et al., 2020). This approach provides confident molecular formula identification of metabolites as well as structural information, with the expectation that molecular isomers with the greatest structural similarity will yield the highest scoring MS/MS spectral match (Schymanski et al., 2014).

Recent applications of liquid chromatography mass spectrometry-based metabolomics have resolved thousands of distinct chemical components that are released by microbes and microbial communities as DOM (the ‘exometabolome’), enabling studies that compare their relative concentrations across environmentally relevant organisms (e.g., Becker et al., 2014; Shibl et al., 2020; Brisson et al., 2021; Ferrer-González et al., 2021). However, only a small fraction of the metabolome is typically identified, with the compounds available in spectral databases matching to only 10s to 100s of features out of thousands of those detected. (e.g., Jiménez-Sánchez et al., 2015; Folberth et al., 2020; Charpentier et al., 2022; Wang and Liu, 2023). Though in silico fragmentation and molecular networking approaches (e.g., Dührkop et al., 2019; Hoffmann et al., 2022; Morehouse et al., 2023) have increased the annotation rate of metabolomics datasets, these tools have additional uncertainties and would also benefit from alternative annotation approaches (Chao et al., 2020).

As the leading tool for characterizing the molecular composition of dissolved organic matter, ultrahigh resolution FT-ICR mass spectrometry provides a means to identify a greater proportion of microbial exometabolomes (Hendrickson et al., 2015; Shaw et al., 2016). Today’s highest resolving mass analyzer, a hybrid linear ion trap/21 Tesla FT-ICR MS that enables automatic gain control (Page et al., 2005), routinely assigns tens of thousands of elemental compositions to marine DOM compounds at sub-ppm mass accuracy with resolving powers > 3,000,000 at *m/z* 200. This custom-built instrument is capable of confidently assigning molecular formulae to complex organic mixtures (e.g. marine DOM, weathered oil) on chromatographic timescales (Smith et al., 2018). These analyses are enabled by advanced data processing tools such as CoreMS (Corilo et al., 2021), a Python-based platform that automates and curates signal processing, calibration, and annotation of mass spectrometry data. However, the cost and accessibility of state-of-the-art LC-FT ICR MS analyses is a significant barrier to conducting large scale metabolomic analyses on such platforms, compared to lower resolution commercially available Orbitrap or time-of-flight instruments that are most used in metabolomic studies.

Here, to address the low annotation rates of tandem MS approaches for metabolomic studies, we developed a novel pipeline that vastly increases the rate of metabolite annotation in environmental samples. Our approach couples the unparalleled resolving power and mass accuracy of the 21T FT-ICR-MS, which we use to generate a library of molecular formulas in small subset of samples, with the relative accessibility of Orbitrap instruments, which we use to detect and identify metabolites in the library across environmental gradients. Similar approaches have been previously employed to increase the confidence and throughput of proteome measurements (Smith et al., 2002).

We used our new approach to determine the effects of iron (Fe) deficiency on the exometabolome of *Phaeodactylum tricornutum* (*P. tricornutum)*, a well-studied genetic model for marine diatoms and biofuel algae (Martino et al., 2007; Bowler et al., 2008; Butler et al., 2020; Song et al., 2020). Previous studies found major transcriptional shifts in *P. tricornutum* in response to Fe deficiency, a commonly encountered nutrient stress in the ocean, but the influence on *P. tricornutum’s* exometabolome remain largely unknown. Recent LC-MS metabolomic analyses of *P. tricornutum* identified 26 exometabolites and demonstrated how algal exometabolite composition may influence phycosphere bacteria community structure (Brisson et al. 2023). Measuring exometabolome changes under Fe limitation can reveal algal adaptation to stress and its impact on microbial interactions, but requires more complete metabolite annotations. Our results (1) demonstrate how FT-ICR MS formula library annotations complements MS/MS based annotations with added confidence and coverage, and (2) reveal specific changes in *P. tricornutum’s* exometabolome in response to Fe stress, such as decreased lipid production and shifts in protein and phytochemical composition. The formula library annotation strategy is designed to be openly accessible and flexible, enabling its adaptation to other metabolomic studies that would benefit from confident molecular formula annotations.

## 2. Material and Methods

### 2.1. Culturing and Sample Generation

Cultures of the diatom (*P. tricornutum* Bohlin strain CCMP 2561) were maintained in modified f/2 medium (Guillard and Ryther, 1962; Smith and Chanley, 1975), with Fe added to a final concentration of 1 μM. Cultures were maintained at 19.0 ± 0.5°C under cool white, fluorescent lights (2030 lux) on a 12-hour cycle. Parent culture density was monitored by manual cell counts using a hemocytometer, as well as by fluorescence measurements (Spectramax Gemini EM fluorescence microplate reader, λ_Ex_ = 440 nm, λ_Em_ = 680 nm). To pre-treat the diatom for this experiment, the axenic *P. tricornutum* culture was transferred to f/2 with 10 nM dissolved Fe at a cell density of 10^3^ cells/mL. This concentration was selected based on initial experiments comparing *P. tricornutum* growth in full f/2 medium (11.7 µM Fe), with 1 µM or 10 nM added iron, and with no Fe added to the medium. The 10 nM Fe condition limited algal growth, so this was chosen as the “Fe deficient” treatment for these experiments. This is on par with findings in previous work, where Fe limitation of *P. tricornutum* was documented at total Fe concentrations of 5-10 nM (Allen et al., 2008; Castell et al., 2021). There was no observed difference between the full f/2 preparation and the 1 µM Fe treatment, so the 1 µM Fe treatment was selected as the “iron replete” condition. The pre-treated parent cultures were allowed to grow until they reached a cell density of ~10^5^ − 10^6^ cells/mL (Fe limited cultures could not reach the target cell density of 10^6^ cells/mL), then transferred to fresh media to a final concentration of 10^3^ cells/mL.

After three rounds of pre-conditioning in medium containing 10 nM Fe, the diatoms were transferred to the experimental conditions. 100 mL f/2 medium in 250 mL polycarbonate flasks (Triforest) were inoculated from the triple pre-treated low Fe parent for a starting cell density of 1000 cells/mL in triplicate for both the Fe replete and Fe deficient treatments. Cultures were maintained under the same incubation conditions as the parents and monitored by fluorescence measurements. By day 7, growth differences between the Fe replete and Fe deficient cultures emerged, with significantly greater abundance and relative fluorescence in the Fe replete treatment (p value < 0.05, t-test) (Fig. S1). 10 mL subsamples were collected and filtered (0.22 µm, Whatman) after 1, 7, and 16 days of growth (T1, T7, and T16, respectively) for LC-MS metabolomics (Fig. S1). Samples collected at T1 were used to constrain the abundance of exometabolites present within the initial growth media, as cells were still in lag phase and media composition was still reflective of background concentrations. To compare the abundance of exometabolites present in the Fe deficient versus Fe replete conditions, samples were collected when both cultures were in logarithmic growth phase (T16). At T7, Fe replete cultures were in early logarithmic growth phase and had reached a similar cell density to the Fe deficient treatment at T16, and the samples from this time point were used to investigate whether the abundance differences between Fe replete vs. deficient treatments related to differences in growth stage or differences in Fe nutrition.

### 2.2. Sample Preparation

Samples were solid phase extracted using 0.1 g PPL cartridges (Agilent Technologies) following standard methods (e.g., Dittmar et al., 2008; Mitra et al., 2013; Boiteau et al., 2019). Cartridges were primed with 3 mL LC-MS grade methanol (VWR Chemical), 3 mL 0.1% trace metal-grade HCl (Fisher Chemical), and 3 mL ultrapure water. 10 mL samples from each culture were filtered (0.22 um PES, Whatman) and loaded onto the primed columns. Columns were rinsed with 3 mL ultrapure water, and frozen at −20 °C until analysis. To prepare for analysis, samples were eluted in 1.5 mL LC-MS grade methanol, dried down to an approximate volume of 50 μL, and rehydrated with ultrapure water to a final volume of 1 mL. 500 μL aliquots of each sample were transferred to 2 mL polypropylene HPLC vials (VWR) and spiked to 1 μM cyanocobalamin (Sigma Aldrich), which served as an internal standard. Equal volumes of each sample from the same time point were mixed to create pooled samples for quality control.

### 2.3. LC-Orbitrap MS

Chromatographic separation of the sample was performed using a bioinert Dionex Ultimate 3000 LC system. The sample loop was filled with 50 μL of sample, which was pushed onto a Zorbax XDB-C18 3.5 um column by the nanopump at 30 μL/min (95% ultrapure water + 5 mM ammonium formate, 5% LC-MS grade methanol + 5 mM ammonium formate). Samples were separated with a 30-minute gradient from 95% solvent A (ultrapure water + 5 mM ammonium formate) and 5% solvent B (methanol + 5 mM ammonium formate) to 95% solvent B, followed by a 5-minute isocratic elution period at 95% solvent B to flush remaining compounds off the column. Solvent composition was switched back to 95% solvent A, and held there for 9 minutes to re-equilibrate the column before introducing the next sample. All chemicals and solvents were LC-MS Grade (Optima; Fisher Scientific). The column was held at 40°C throughout the run. Column eluent was directed into a Thermo Scientific Orbitrap IQ-X mass spectrometer equipped with a heated electrospray ionization source. ESI source parameters were set to a capillary voltage of 3500 V, sheath, auxiliary and sweep gas flow rates of 5, 2, and 0 (arbitrary units), and ion transfer tube and vaporizer temperatures of 275 °C and 75 °C. MS^1^ scans were collected over a *m/z* range of 100-1000 in high resolution (500k at *m/z* 200, transient length 1024 ms) positive mode. MS^2^ spectra were collected in data-dependent acquisition mode in the ion trap, with an HCD energy of 30 and a MS^2^ isolation width of 1.60.

### 2.4. LC-21T FT-ICR MS

Chromatographic separation of samples was identical to the method used for data collected using LC-Orbitrap MS. All chemicals and solvents were Honeywell. The column was held at 40 °C throughout the run. The eluent was coupled to a heated electrospray source operated in positive mode (3.75kV) on the custom-built hybrid linear ion trap FT-ICR mass spectrometer equipped with a 21T superconducting solenoid magnet (Hendrickson et al., 2015). The inlet capillary and source heater temperatures were set to 350 °C and 75 °C, respectively. The sheath and auxiliary gas flow rates were set to 5 and 3 (arbitrary units). MS^1^ spectra were collected from *m/z* 200 to 2000 with a target resolution of 1,200,000 at 200 *m/z* (transient length of 1536 ms), an AGC target of 1×10^6^ charges, and a maximum ion injection time of 500 ms.

### 2.5. Feature List Generation and MS/MS Analysis

Individual samples were run by LC-Orbitrap as described above, and processed using MSDial (Tsugawa et al., 2015, 2020; ver.5.1.230517) to generate a feature list. Only peaks with signal intensity above 50,000 amplitude were considered in the feature list to exclude analytical noise. Mass accuracy for singly-charged species was constrained by 0.01 Da and 0.1 Da for MS^1^ and MS^2^ spectra, respectively. An identification score cutoff of 70% were imposed to limit false matches to the Pos VS17 authentic standard spectral library (324,191 entries) were accurate. To generate the feature list, features were aligned with a MS^1^ tolerance of 0.01 Da and a retention time tolerance of 1 minute to account for minor matrix effects.

### 2.6. Library Generation and Feature Annotation

A pooled sample was generated from cultured samples (n = 18) and analyzed using LC-21T FT-ICR-MS as described above. The mass spectra were averaged over two-minute time intervals and internally calibrated based on a series of polysiloxane masses that appeared throughout the chromatogram, which improved mass accuracy and reduced processing time compared to annotation of individual spectra. Using the open-source platform CoreMS (Corilo et al., 2021), molecular formulae were assigned using stoichiometric ranges that encompass most biologically relevant compound classes (e.g., Stubbins et al., 2010; Rivas-Ubach et al., 2018a): C 1-50, H 4-100, O 1-16, N 0-8, S 0-2, P 0-1. Additional constraints of a max DBE of 18, maximum O/C of 1.2, and maximum H/C of 3 were imposed on formula assignments to ensure resulting formulas were reasonable. 69.5% of the features found in the pooled sample were assigned a molecular formula. These formulas were then compiled into a library, which was then used to annotate features present in the metabolomics feature list based on matching the mass and retention time of the features in the feature list and molecular formula library.

## 3. Results and Discussion

### 3.1. Exometabolome annotation using LC-MS/MS and standard reference database

Metabolites detected by LC-Orbitrap MS were annotated across the samples using MSDial. A total of 5,403 metabolite features (*m/z* detected at a specific retention time) were compiled. Annotation of these metabolites based on comparison to the MS/MS spectra of 324,191 authentic standards yielded 318 assignments, or an annotation rate of 5.9%. This low annotation rate reflects the need for alternative approaches that yield more comprehensive coverage and motivated the development of the ultrahigh resolution mass spectrometry molecular formula library-based annotation approach described in this manuscript.

The MS/MS annotation strategy was primarily limited by the number of metabolites in spectral libraries, which are focused on small molecules that are commercially available or present in well studied model organisms. Most of the MS/MS annotations were of low molecular weight (*m/z* < 300) features, though some features up to *m/z* 430 were annotated (Fig. 4). 94% of *P. tricornutum* exometabolites’ MS/MS spectra did not match any of the 324,191 database spectra, highlighting a major information gap associated with this approach. We note that these libraries continue to grow, and that there are several emerging in silico tools that can predict fragmentation spectra of molecules (e.g., Dührkop et al., 2019; Hoffmann et al., 2022; Morehouse et al., 2023). However, in silico tools continue to have significant uncertainty associated with them (Chao et al., 2020) and the size of the putative metabolome far exceeds - by orders of magnitude - the size of any current or future practical experimental database. A second limitation of MS/MS annotation was that fragmentation spectra were only measured for a subset of the most abundant detected features (5,312 out of 5,403 total features; Table 1; Fig. 4). Furthermore, some spectra may contain noise or isobaric interferences that resulted in false negative spectral matches, although the frequency of these issues is unclear.

**Table 1:** Number of features annotated and not annotated by the application of the FT-ICR MS Library approach and a MS/MS library approach using MSDial. Fe replete and Fe deficient features were present in the corresponding treatments at T16 at an abundance >2fold greater than T1. “Overlapping” features refer to those present in both the high and low Fe treatments. “Significant” features are those that were expressed significantly differently (>2-fold intensity difference, p < 0.05) between the Fe deficient and Fe replete treatments.

|  | All<br>exometabolome<br>features | Background<br>removed<br>features | Fe<br>replete<br>features | Fe<br>deficient<br>features | Overlapping<br>Fe<br>features | Significant<br>features |
| --- | --- | --- | --- | --- | --- | --- |
| Total | 5403 | 4031 | 3444 | 3003 | 2670 | 668 |
| MS/MS<br>assigned | 318 | 212 | 181 | 162 | 148 | 61 |
| Library<br>assigned | 2840 | 2255 | 1953 | 1772 | 1585 | 450 |
| Outside library<br>$m/z$ range | 1433 | 955 | 807 | 635 | 563 | 142 |
| Inside library<br>$m/z$ range but<br>not detected | 834 | 600 | 515 | 446 | 401 | 51 |
| Detected in<br>library but not<br>assigned | 296 | 221 | 169 | 150 | 121 | 25 |

### 3.2. LC-21T FT-ICR MS library generation & exometabolome annotation

This study introduces an advanced data informatics software that annotates LC-MS exometabolomes (collected on lower resolution instruments such as Orbitrap or time-of-flight) with confident molecular formula assignment based on paired data from ultrahigh resolution 21T FT-ICR MS. Because this instrument is broadly accessible through user facilities but instrument time can be limited by high demand, the approach is designed to analyze a single or small number of representative ‘reference’ sample(s), such as a pooled sample containing a mixture of samples across all treatments and replicates.

Molecular formulas were assigned to features in the LC-FT-ICR MS raw data using CoreMS, which were then compiled into a library of formulas (Fig. 1). 69.5% of the 57,512 detected mass peaks in the FT-ICR MS data were annotated with molecular formula within a mass error of 0.3 ppm. This level of mass accuracy is an order of magnitude better than that achieved by the Orbitrap (Fig.3), highlighting the power of high field ICR instruments when exact mass measurements are crucial for analyses.

**Fig. 1:**
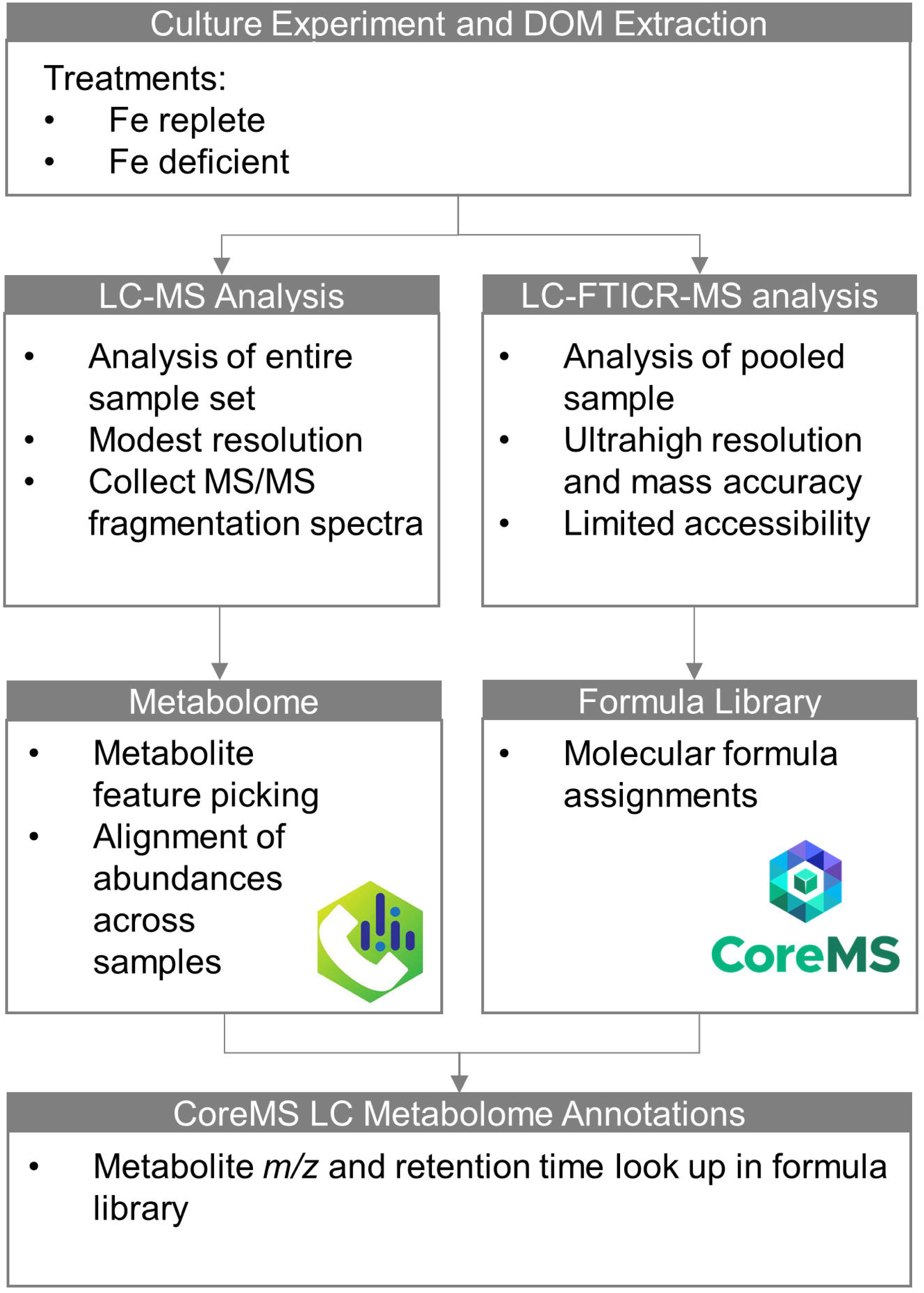
Overview of CoreMS metabolomics formula assignment workflow. All samples were analyzed by LC-Orbitrap MS, and data was processed in MSDial to yield a feature list with 5403 features. Only 318 had annotations based on MS/MS similarity scoring (5312 of the features had collected MS/MS spectra), highlighting a limitation of this annotation approach. A pooled sample was analyzed with the same chromatography coupled to 21T FT-ICR MS. The mass spectra were binned in 2-minute time intervals and recalibrated based on a set of polysiloxanes observed across the chromatography. Molecular formula assignments were made based on elemental selection criteria described in the methods. This library was used to annotate molecular features in the metabolomics feature list based on matching the mass and retention time of the features in the metabolomic data set and the FT-ICR MS molecular formula library.

The LC-Orbitrap metabolome was then annotated with molecular formula assigned in the FT-ICR-MS library (Fig. 1), accounting for a minor retention time shift between instruments (Fig. S2). A library match was defined as an FT-ICR MS mass matching the metabolite mass within 3 ppm (selected based on the mass accuracy and resolving power of the Orbitrap MS used in this study) that occurred within the same 2-minute retention time window. Of the 5,403 exometabolome features, 3,175 had library matches, and 39 of these had two library matches – these multiple library matches indicate features that were not fully resolved in the lower resolution mass spectra. This highlights another advantage of the ultra-high resolution library approach; even the highest resolution mode of the Orbitrap MS still fails to resolve some compounds of similar molecular weights in complex DOM (Olsen et al., 2005; Makarov et al., 2006; Scigelova and Makarov, 2006), and these metabolites are flagged by the CoreMS LC exometabolome annotation pipeline. In the event of multiple library hits, the molecular formula of the library feature with the highest signal to noise ratio was reported, but ultimately these features should be considered with caution as they may represent multiple distinct molecules.

To gain additional insight into *P. tricornutum*’s annotated exometabolome, the molecular formulas were classified based on stoichiometric elemental ratios (Fig. 2) following previously established criteria (Rivas-Ubach et al., 2018a). The molecular formulas identified in this analysis were classified as a wide range of biomolecules, encompassing peptides, lipids, carbohydrates, phytochemicals, sulfur-containing organics, and phospholipids. While this is a useful framework for assessing the composition of *P. tricornutum*’s exometabolome, it is worth noting that the stoichiometric classifications are accurate predictions for most, but not all biochemicals, due to some overlap in the stoichiometric ratios of different categories (Rivas-Ubach et al., 2018a). Most of the molecular formulas assigned using the pipeline fell into the lipid, peptide, or phytochemical categories. As *P. tricornutum* is a model species for biofuel production due to its high lipid content, the dominance of lipids in its exometabolome in this experiment was unsurprising (Chauton et al., 2013; Vandamme et al., 2018). Furthermore, the abundance of peptides is also consistent with the high protein content of *P. tricornutum*, which can account for up to 25% of dry weight (Song et al., 2020). Diatoms are also known to produce a range of polyphenolic phytochemicals, and their production is in part regulated by metal stress (Rico et al., 2013). Our focus on these classes is also due in part to the methodological approach employed; solid phase extraction retains these classes well (Dittmar et al., 2008; Miranda et al., 2023), and reverse-phase liquid chromatography with positive mode electrospray ionization mass spectrometry separates and detects these classes effectively.

**Fig. 2:**
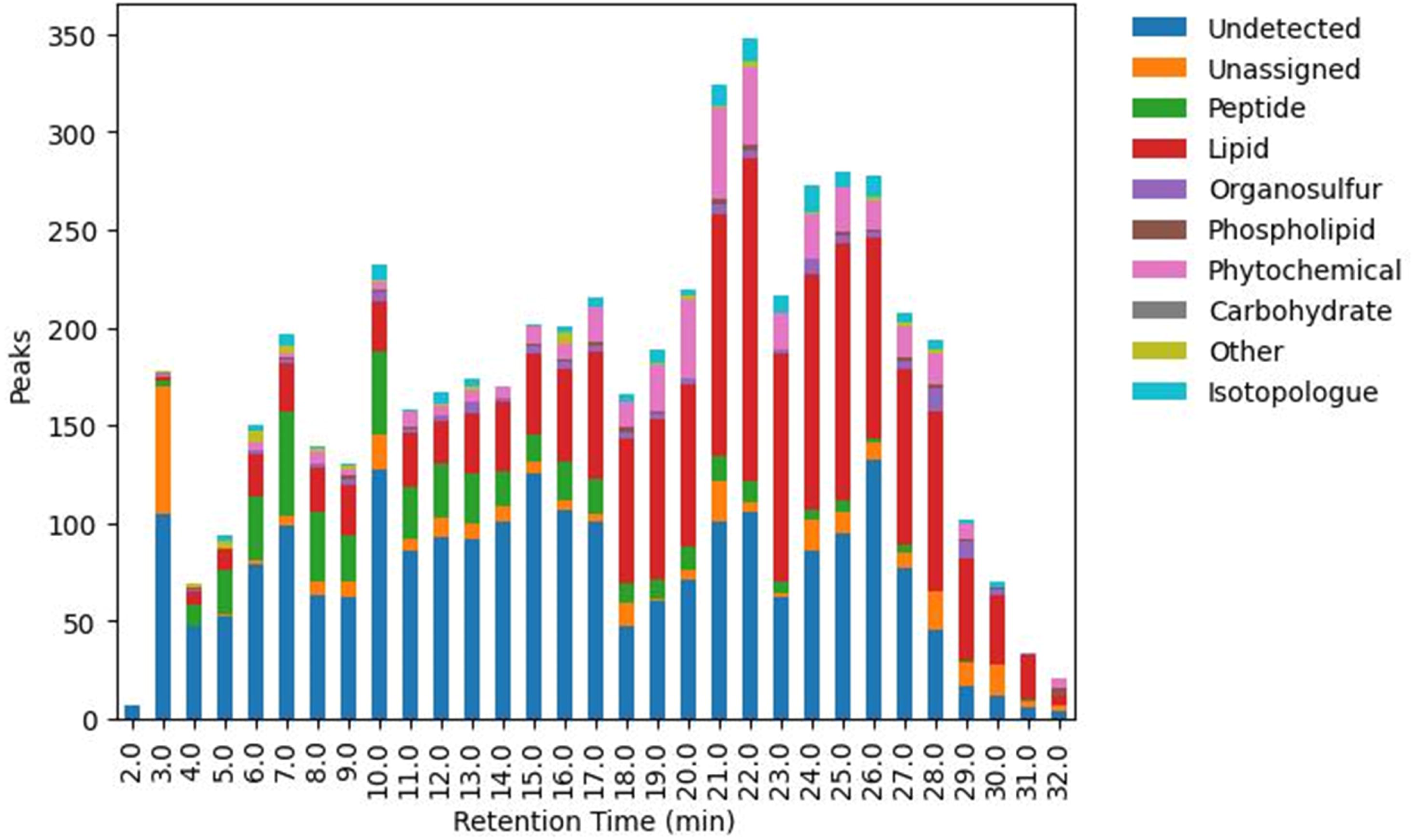
Stoichiometric classifications of exometabolome features annotated with the formula library approach. Classification categories were based on elemental ratios of molecular formula adapted from (Rivas-Ubach et al., 2018a). The ‘Undetected’ category corresponds to metabolite features that were not detected by LC-ICR MS. The ‘Unassigned’ category corresponds to metabolite features that were detected by LC-ICR MS but not assigned molecular formula.

Of the library matches, 2,840 of these were assigned molecular formulae, representing an annotation rate of 53.2% of the exometabolome features. The quality of the molecular formula assignments was assessed by the distributions of mass errors (Fig. 3). The mass errors of the FT-ICR MS assignments show a narrow, symmetric, and unimodal distribution centered around zero (Fig. 3), as expected for well calibrated and accurately assigned data. The mass errors of the average exometabolome masses measured by Orbitrap MS also show a symmetric and unimodal distribution, but with much less mass precision and accuracy. Since the reliability of molecular formula assignment is directly related to mass accuracy, this comparison demonstrates the utility of using ultra-high resolution and accurate mass data to annotate lower-resolution metabolomes.

**Fig. 3:**
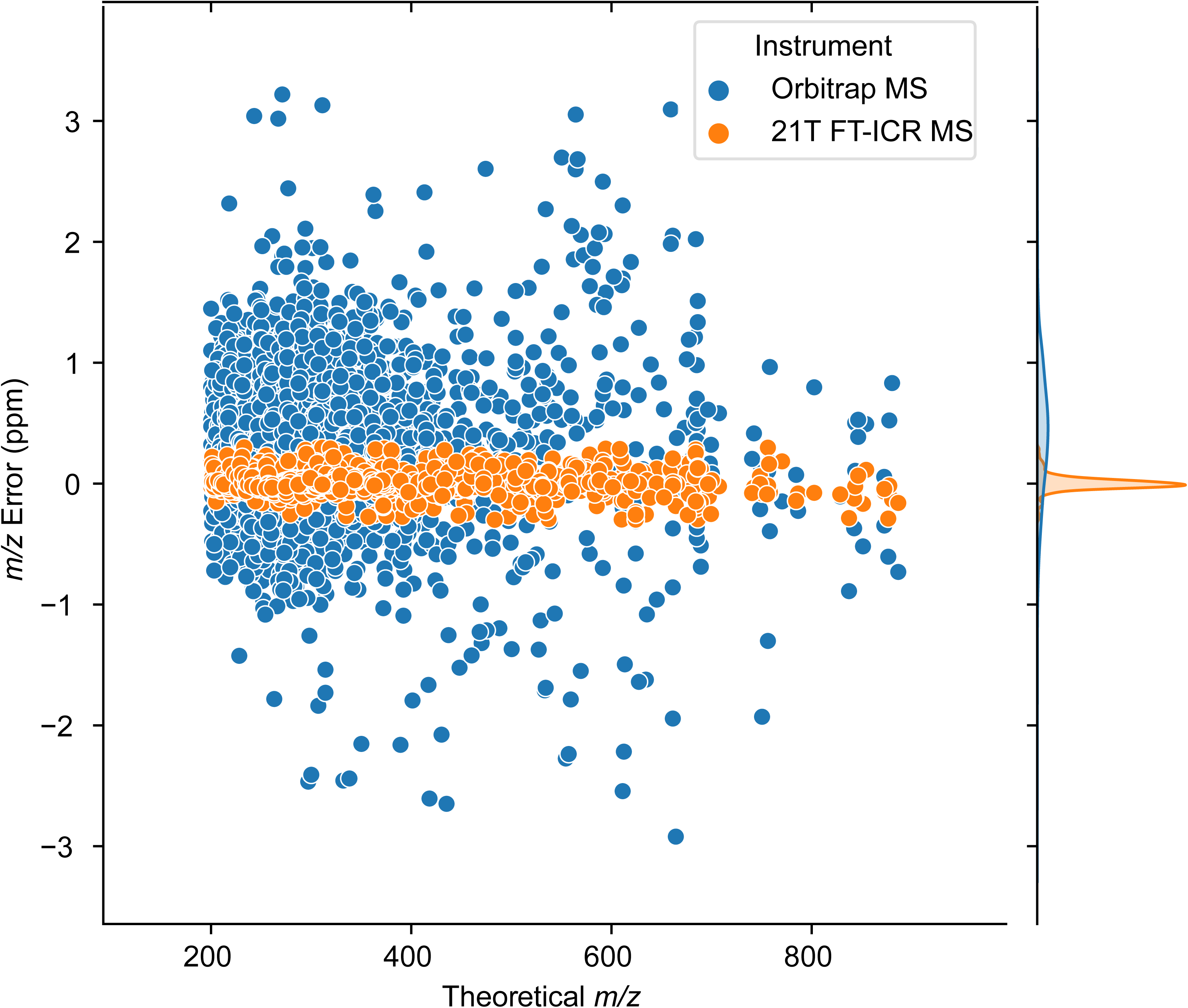
Scatter plot showing the mass error of FT-ICR MS formula library annotated metabolites. Mass error was calculated as the difference between the average *m/z* measured by LC-Orbitrap or LC-ICR MS and the theoretical ion mass, with kernel density estimate distribution at the top. The 10-fold greater mass precision of the 21T ICR-MS measurements highlights the advantage of this instrumentation for assigning accurate formula.

### 3.3. Comparison of MS/MS and LC-21T Library Annotations

To evaluate the strengths and limitations of each strategy, we next compared the annotations of the FT-ICR MS library approach to the MS/MS fragmentation library approach. A total of 166 metabolites were annotated with both approaches, and 94% of the assigned molecular formula agreed between the two methods, providing added confidence in our library annotations. In the case of the 10 mis-matched formula, the mass error of the MS/MS assigned formula were consistently outside of the expected mass accuracy of the Orbitrap MS (> 3 ppm error), except for one molecular formula that was annotated as an isotopologue of another peak by the FT-ICR MS library. These results indicate that the FT-ICR MS formula assignments were more robust. This analysis also demonstrates that MS/MS spectra matching does not always yield correct molecular formula annotations, even when conservative criteria for mass tolerance and spectral similarity score are used. The use of stricter criteria reduced the number of incorrect matches, but at the expense of also reducing the number of correct matches and thus overall annotation rate. Despite these findings, a major benefit of the MS/MS fragmentation approach is that the fragment analysis provides additional information on the putative molecular structure rather than just a formula. These two approaches could be used complementarily, with the structural information from the MS/MS match informing on the confident formula assignment from the library approach. However, as the rates of MS/MS annotation are still low, comprehensive coverage of features is still an area to be improved.

Though the annotation rate of the molecular formula library approach was considerably higher than that achieved by the MS/MS matching approach, it still left 46% of the exometabolome unannotated. 59% of these unannotated metabolites were smaller than *m/z* 200 and thus simply below the minimum detectable mass of the 21T FT-ICR MS. 33% of the unannotated features were not detected despite being inside of the FT-ICR MS mass range. This may have been due to differences in the sensitivity of the instruments or because many compounds were diluted in the pooled samples relative to individual samples collected from later time points where higher abundances were generally observed. Finally, only 296 features were detected by the 21T FT-ICR MS and not assigned a molecular formula (Table 1) – 5.5% of the entire dataset. The percent of detected but unassigned features were similar across the *m/z* range (Fig. 4). These metabolites likely include elements that were not considered in the formula assignment criteria used for this study. Of the 3,136 exometabolome features detected in the pooled sample by FT-ICR MS at 21T, the molecular formula library pipeline achieved a 90.5% annotation rate. These results emphasize the potential of ultra-high resolution mass spectrometry to greatly improve the coverage of LC-MS exometabolome annotations.

**Fig. 4:**
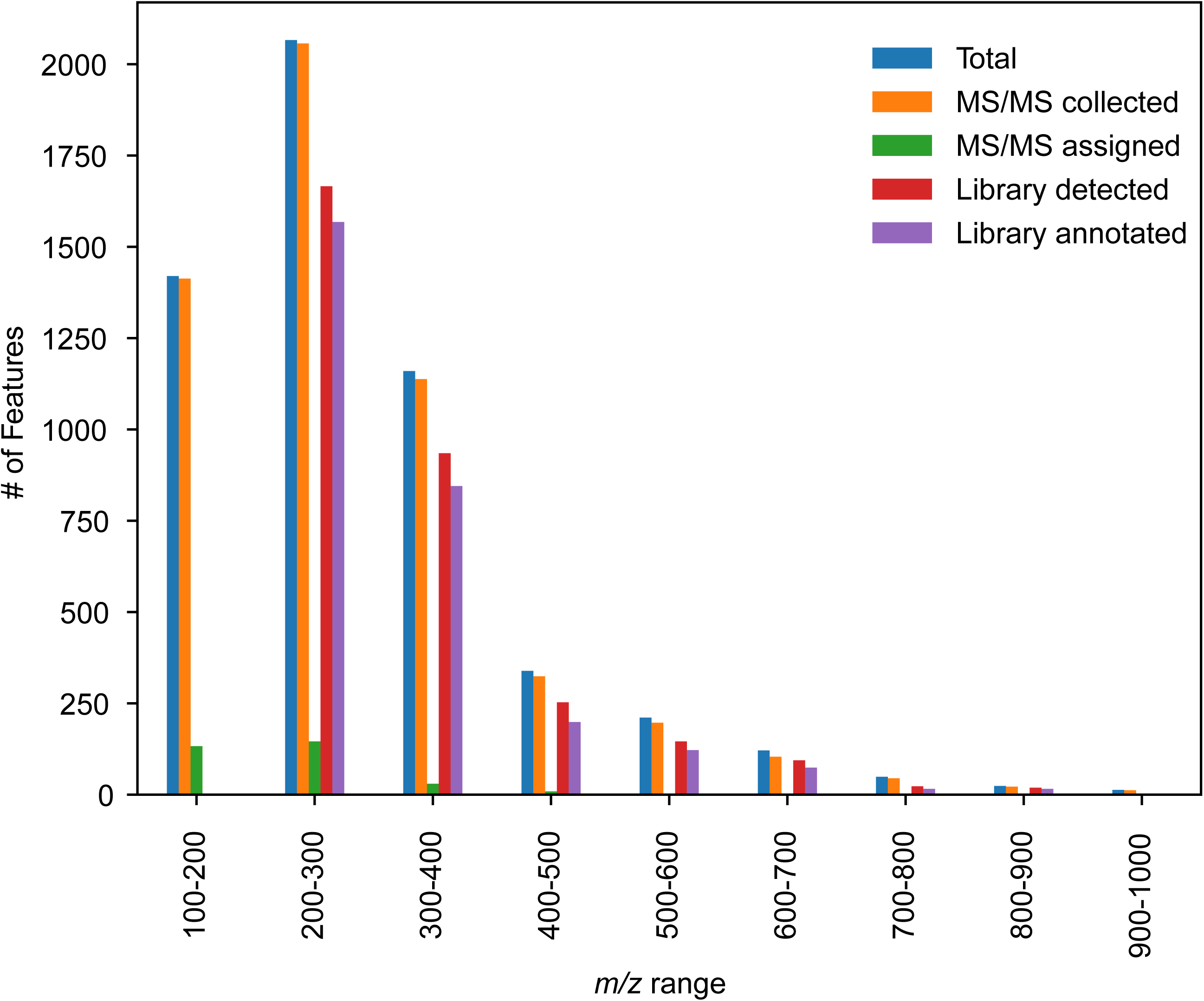
*m/z* distribution of the total number of identified features (blue), features with collected MS/MS fragmentation spectra (ion trap of the Orbitrap IQ-X; orange), features assigned by a conventional MS/MS matching approach using MSDial (green), and features detected (red) and assigned (purple) using the 21T FT-ICR MS library annotation approach developed in this study. Despite the high availability of MS/MS spectra for metabolite features, less than 10% were annotated by matching to the MSDial authentic standard spectral library (Pos VS17). Features below *m/z* 200 were below the mass range of the FT-ICR-MS formula library and were thus not annotated with this approach.

### 3.4. Differentially Expressed Metabolites in High vs Low Iron Conditions

With the development of this new annotation pipeline, we were able to evaluate changes in *P. tricornutum*’s exometabolome between Fe replete and Fe deficient growth conditions. First, we filtered out 1372 background features that did not exhibit at least 2-fold greater maximum abundance in the late log samples relative to the average of the initial timepoint samples. Of the remaining features, 3444 metabolites accumulated in the culture media under Fe replete conditions (2-fold greater average abundance at late-log versus initial timepoints), while 3003 accumulated under Fe deficient conditions. Many of these metabolites accumulated in both treatments (2670), indicating that they are generally released regardless of Fe condition.

To investigate metabolites that reflected the Fe stress state of the diatoms, we identified features that were significantly different in abundance between treatments in the late-log samples (>2-fold difference between treatments, Bonferroni adjusted two-tailed t-test p-value <0.05), which yielded a list of 668 features. Of these features, MS/MS matching annotated 61 features, while the molecular formula library annotated 450 (Table 1, Fig. 5). Most of the statistically significant features were more abundant in the Fe replete condition (Fig. 5b). Since the abundances of most of these metabolites also increased in the Fe deficient cultures relative to the initial time point (Fig. S5), it is likely that their abundance reflects the cell density in the culture rather than the direct effect of Fe stress. In contrast, the 25 metabolites that were in greater abundance in the Fe deficient treatment were mostly absent from the Fe replete treatment, both at the late-log time point as well as the earlier time point when the Fe replete culture had similar cell density to the Fe deficient culture (T7; Fig. S6). Thus, these exometabolites appear to reflect a metabolic response to the Fe deficient culture conditions. Although MS/MS matching was not able to annotate any of these features, eleven were annotated using the FT-ICR MS library approach with *m/z* errors below 0.2 ppm (Fig 6c; Table 2). This highlights the advantage of the library annotation pipeline for investigating metabolites that are not well-represented in MS/MS fragmentation databases.

**Fig. 5:**
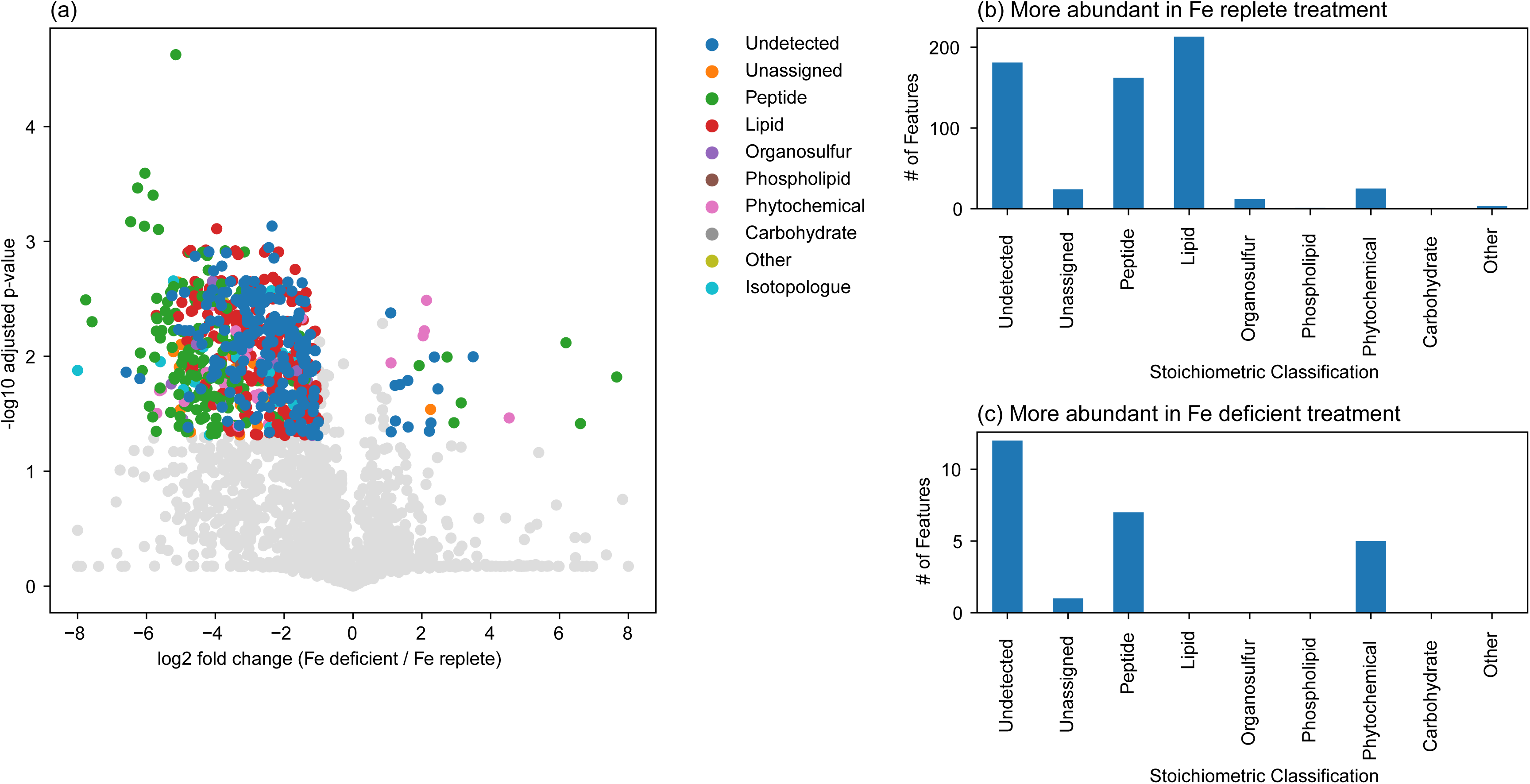
(a) Volcano plot of metabolites, showing the stoichiometric classification of statistically significant features (adjusted p-value < 0.05) that differ (> 2-fold) in relative abundance between the Fe deficient versus Fe replete treatments on Day 16. (b-c) Bar charts showing the number of significantly different features that were more abundant in the Fe replete (b) or Fe deficient treatments (c) belonging to each stoichiometric class. Annotated features in panel (c) are listed in Table 2.

**Table 2:** Annotated features present at higher abundances in the Fe-deficient condition. These features all fall within stoichiometric classifications of peptides or phytochemicals, likely reflecting changes in chlorophyll biosynthesis under Fe limitation (e.g., Allen et al., 2008; Zhao et al., 2018).

| Molecular Formula | Average $m/z$ [M+H]<sup>+</sup> | Retention Time (min.) | FT-ICR MS Library $m/z$ Error (ppm) | Stoichiometric Classification | Log 2-fold difference (Fe deficient/ Fe replete) | Adjusted p-value |
| --- | --- | --- | --- | --- | --- | --- |
| C<sub>13</sub> H<sub>14</sub> O<sub>3</sub> | 219.1015 | 15.3 | +0.072 | Phytochemical | 2.04 | 0.0066 |
| C<sub>14</sub> H<sub>14</sub> O<sub>3</sub> | 231.1017 | 12.8 | -0.010 | Phytochemical | 2.14 | 0.0032 |
| C<sub>14</sub> H<sub>16</sub> O<sub>4</sub> | 249.1124 | 12.9 | -0.026 | Phytochemical | 2.07 | 0.0060 |
| C<sub>11</sub> H<sub>16</sub> O<sub>3</sub> N<sub>4</sub> | 253.1297 | 7.6 | -0.034 | Peptide | 1.92 | 0.012 |
| C<sub>13</sub> H<sub>14</sub> O<sub>4</sub> S<sub>1</sub> | 267.0687 | 16.7 | +0.049 | Phytochemical | 1.10 | 0.011 |
| C<sub>15</sub> H<sub>28</sub> O<sub>2</sub> N<sub>2</sub> | 269.2225 | 19.4 | +0.096 | Peptide | 2.73 | 0.010 |
| C<sub>12</sub> H<sub>12</sub> O<sub>2</sub> N<sub>4</sub> S<sub>1</sub> | 277.0760 | 11.0 | +0.052 | Peptide | 6.61 | 0.038 |
| C<sub>13</sub> H<sub>20</sub> O<sub>3</sub> N<sub>4</sub> | 281.1610 | 9.1 | +0.041 | Peptide | 2.93 | 0.038 |
| C<sub>13</sub> H<sub>17</sub> O<sub>2</sub> N<sub>5</sub> S<sub>1</sub> | 308.1179 | 8.3 | -0.013 | Peptide | 6.19 | 0.0076 |
| C<sub>12</sub> H<sub>12</sub> O<sub>4</sub> N<sub>4</sub> S<sub>1</sub> | 309.0655 | 9.8 | +0.036 | Peptide | 7.66 | 0.015 |
| C<sub>14</sub> H<sub>34</sub> O<sub>10</sub> N<sub>5</sub> Na<sub>1</sub> | 456.2277 | 12.9 | -0.089 | Peptide | 3.14 | 0.025 |

The molecular annotations provide insight into the specific effect of Fe stress on exometabolite composition. Peptide and phytochemical-classified compounds exhibit greater shifts in response to Fe deficiency, where some molecules became more abundant while others less abundant. Here, phytochemicals are defined as oxy-aromatic compounds as per constraints proposed by Rivas-Ubach et al. 2018a and encompasses compounds such as flavonoids, lignin, and chlorophylls. Previous work with *P. tricornutum* and other marine diatoms has documented the down-regulation of genes related to processes with high Fe requirements such as nitrate reduction, as well as shifts in expression of different photosynthetic components (Allen et al., 2008; Lommer et al., 2012; Marchetti et al., 2012; Zhao et al., 2018). Interestingly, although overall chlorophyll content of Fe-starved diatoms is decreased relative to an iron-replete condition (Greene et al., 1991; Rivas-Ubach et al., 2018b), some genes involved in chlorophyll biosynthesis are up-regulated in response to Fe limitation (Allen et al., 2008). These observations may relate to the specific phytochemicals and peptides that were found to increase in relative abundance in the iron-deficient cultures (Fig. 5c). The three molecules that exhibited the largest fold increase in the Fe deficient conditions had very similar molecular formula, C_12_H_12_O_4_N_4_S_1_, C_12_H_12_O_2_N_4_S_1_, and C_12_H_17_O_2_N_5_S_1_, suggesting that they belong to the same class of metabolites. Further work is needed to elucidate the chemical structures of these metabolites and determine their physiological roles. Nevertheless, this analysis demonstrates the utility of FT-ICR MS annotations for providing molecular insight into a new group of *P. tricornutum* exometabolites that relate to Fe stress.

In contrast to the diatom’s mixed response with regard to proteins and phytochemical compounds, far fewer lipid-classified molecules accumulated in the Fe limited treatment compared to the Fe replete treatment. This is consistent with previous observations of a significant decline in *P. tricornutum*’s total fatty acid content under Fe limitation (Wang et al., 2023), and appear to reflect a general downregulation of lipid biosynthesis, though this does not seem to be universal for all algae. Lipid production is stimulated by only slight Fe supplementation in *Dunaliella tertiolecta* (Rizwan et al., 2017), and under low Fe conditions in *Nannochloropsis oculata* (Sabzi et al., 2021). However, the decrease in lipid content in response to Fe stress has been observed for other algal species, such as *Chlorella vulgaris* and *Chlorococcum oleofaciens* (Liu et al., 2008; Rajabi Islami and Assareh, 2020). Our results indicate that stress induced by Fe limitation results in a decrease in lipid production by *P. tricornutum*, emphasizing that algal species have unique lipid production responses to Fe stress, which must be considered when optimizing growth conditions for specific purposes, such as biofuel production.

## 4. Conclusions

The CoreMS data processing pipeline developed in this study provides an effective strategy to extend LC-MS metabolome annotations based on molecular formula libraries generated by ultrahigh resolution FT-ICR MS. In the current study, the FT-ICR MS library approach annotated ten times the number of features in marine algal exometabolomes relative to MS/MS spectra matching, greatly expanding our window into organisms’ molecular responses to their environment. Our findings revealed differences in the release of lipids, peptides, and phytochemicals produced as the diatom experienced Fe limitation. These molecular formula assignments are also useful stepping-stones to structural elucidation, and they enable future targeted studies that assess mechanisms of adaptation or algal-bacterial interactions in response to Fe stress.

Just as databases of MS/MS spectra can be applied to many studies, the LC 21T FT-ICR MS molecular formula library generated in this study can be used as a resource for other LC-MS metabolome studies of *Phaeodactylum Tricornutum*. Both the CoreMS data processing pipeline and the molecular formula library are publicly available to enable their application to other LC-MS metabolome data sets that use similar chromatographic conditions. Furthermore, analyses of other exometabolome and DOM ‘reference samples’ by LC 21T-FT-ICR MS can be used to generate similar molecular formula libraries for other organisms or environments. Ultimately, this approach to increase annotation rates of metabolomic data collected on widely available instrumentation can expedite the discovery of processes that govern the release, transformations, and biological/environmental effects of exometabolites and other molecules that comprise DOM.

## 5. Data Availability

MS data is available in the MassIVE repository under accession #MSV000094385. CoreMS LC Metabolome scripts are available to download from Github at https://github.com/boiteaulab/CoreMS-LC-Metabolome-PT.

## Acknowledgements

This this research was funded by U.S. Department of Energy (DOE) Office of Biological and Environmental Research, SCW1039. The work at LLNL was performed under DOE auspices of the by Lawrence Livermore National Laboratory under Contract DE-AC52-07NA27344. A portion of this work was performed at the National High Magnetic Field Laboratory, supported by the MagLab User Collaboration Grants Program, the National Science Foundation Cooperative Agreement No. DMR-1644779 and the State of Florida. A portion of this research was performed on project awards 10.46936/intm.proj.2021.60081/60000413 and 10.46936/intm.proj.2019.51140/60006695 from the Environmental Molecular Sciences Laboratory, a DOE Office of Science User Facility sponsored by the Biological and Environmental Research program under Contract No. DE-AC05-76RL01830. The authors would like to thank Kathleen Kouba for assistance and support with algal culture maintenance.

