## Supplemental Information for "Annotation of DOM metabolomes with an ultrahigh resolution mass spectrometry molecular formula library"

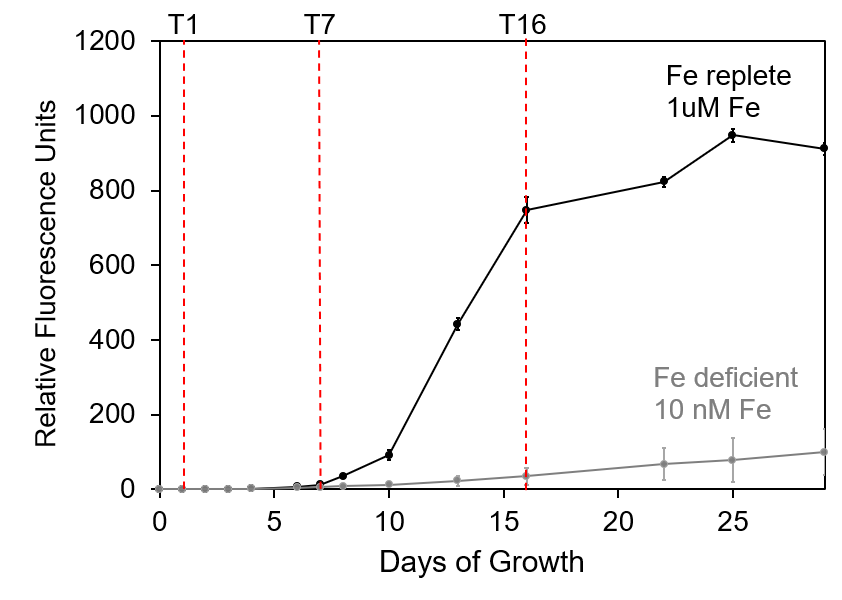

Figure S1: Growth of *Phaeodactylum tricornutum* under iron replete (1uM) and deficient (10 nM) conditions. Cultures were grown in triplicate after 3 rounds of acclimation to iron deficient conditions and started at the same cell density (1000 cells/mL). Subsamples were collected at three timepoints (T1, T7, T16, red lines) and analyzed by LC-MS.

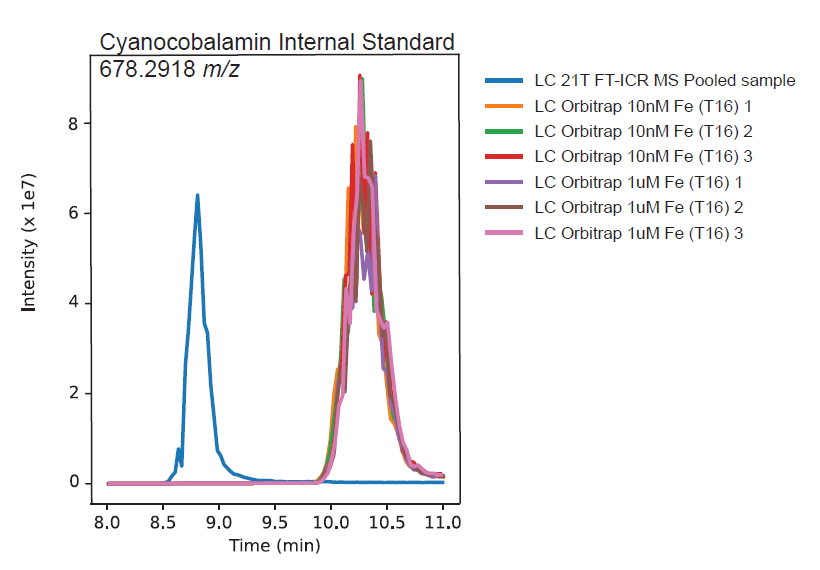

Figure S2: Extracted ion chromatogram of cyanocobalamin internal standard. The 1.43 minute offset between the 21T and Orbitrap data was taken into account during the CoreMS metabolome annotation.

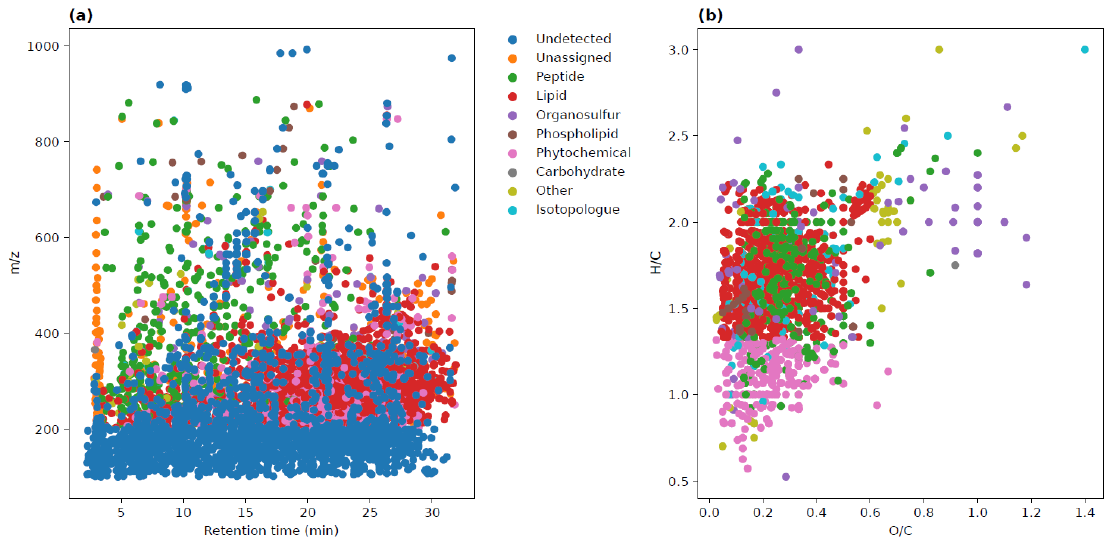

Figure S3: a) Scatter plot of library annotated metabolites, showing the stoichiometric classification (After Rivas-Ubach et al. 2018a) of assigned molecular formula in *P. tricornutum*’s metabolome across the chromatographic separation. The ‘Undetected’ category corresponds to metabolites that were not detected by LC-ICR MS and the ‘Unassigned’ category corresponds to molecules that were detected by LC-ICR MS but not assigned molecular formula. b) Scatter plot showing O/C and H/C ratios of library annotated metabolites.

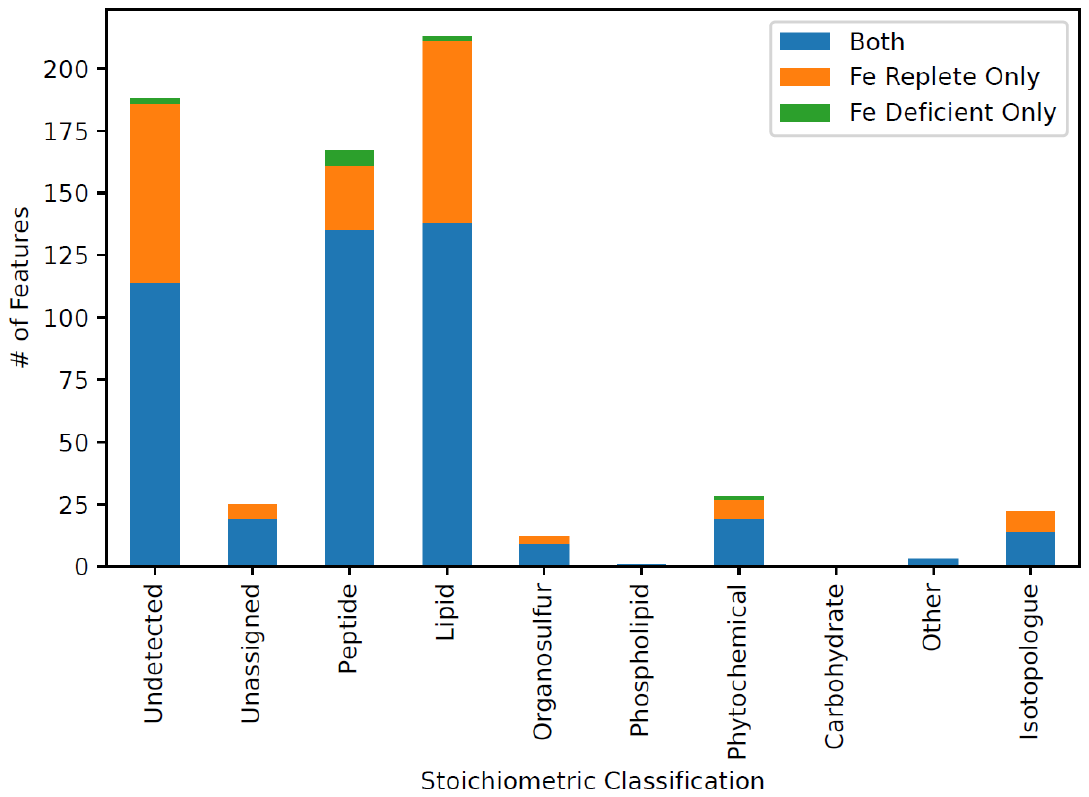

Figure S4: Bar chart showing the number of features within each stoichiometric class that increased in abundance (>2-fold change at T16 relative to T1) in the Fe replete treatment, in the Fe deficient treatment, and in both. These represent the exometabolome of *P. tricornutum* detected by LC-MS.

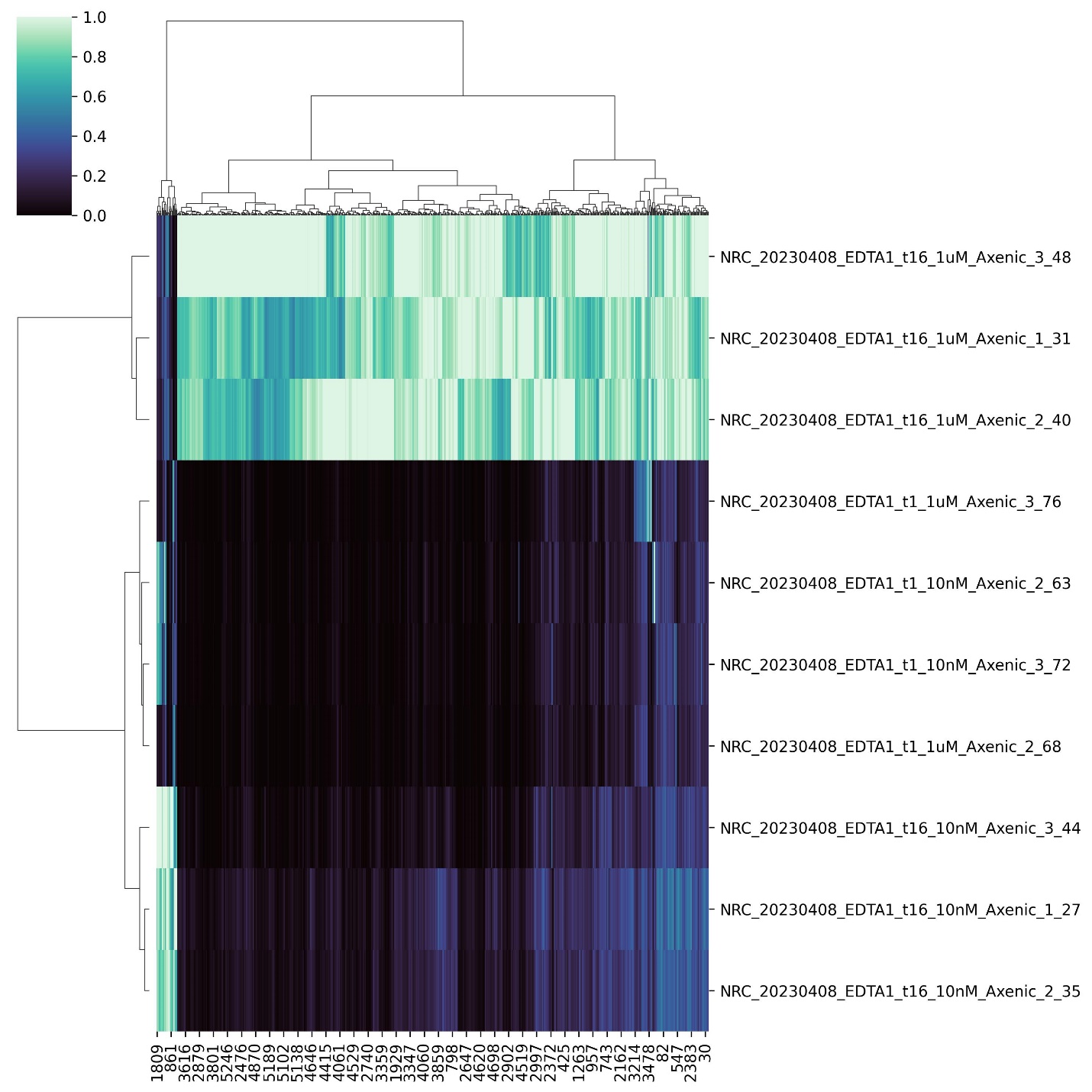

Figure S5: Heat map showing the relative abundance of metabolites that were significantly different between the high and low Fe treatments (adjusted p-value <0.05 and >2-fold change).

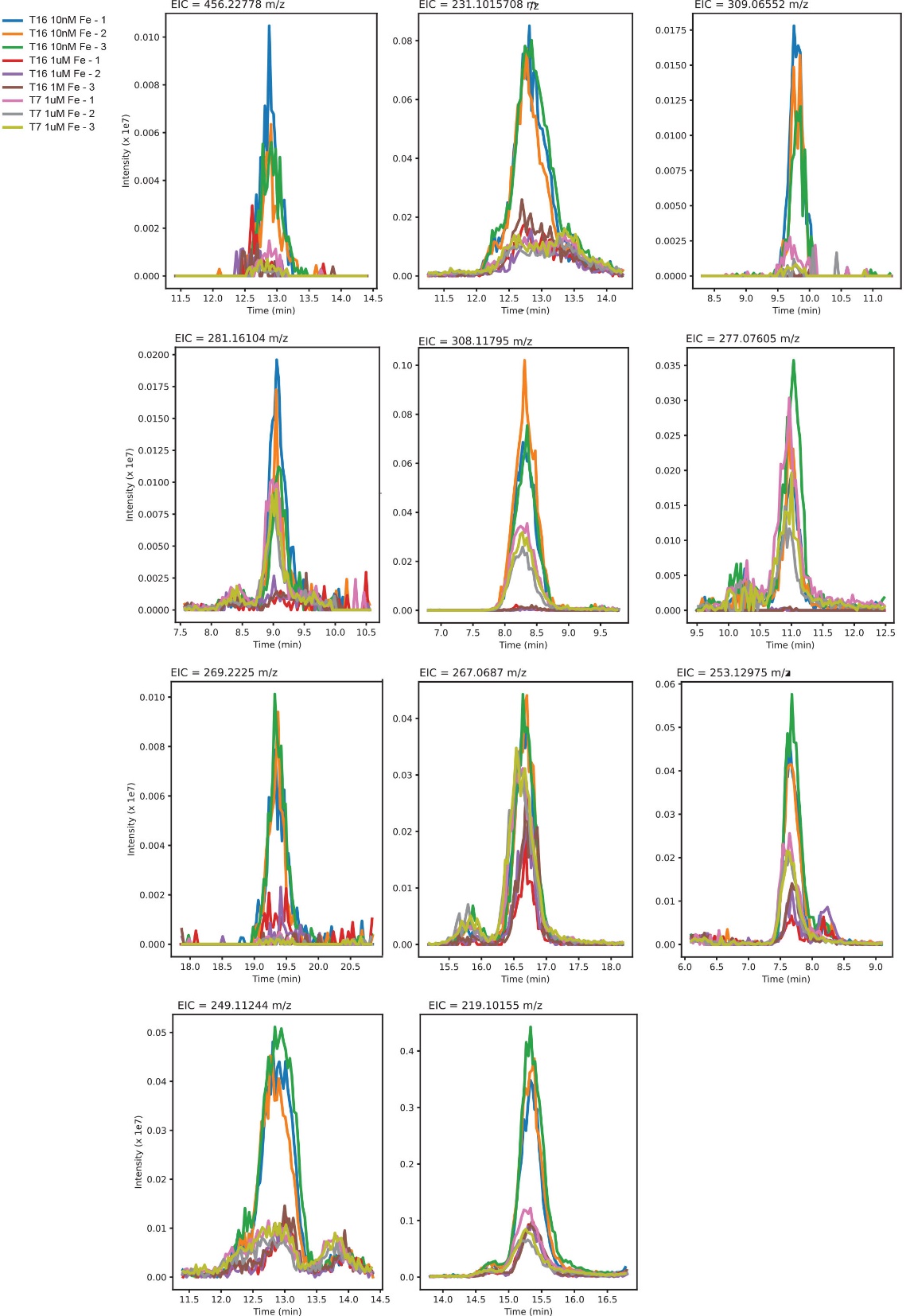

Figure S6:­­­ Extracted ion chromatogram of significant features (adjusted p-value <0.05) that were more abundant (fold change > 2) in the low Fe samples. The overlaid chromatogram includes triplicates from the Fe replete treatments at T7 (early log stage) and T16 (late log stage), comparing them to the Fe deficient treatment at T16 (early log stage).
